# Shifts in the coral microbiome in response to *in situ* experimental deoxygenation

**DOI:** 10.1101/2023.04.06.535971

**Authors:** Rachel D. Howard, Monica Schul, Lucia M. Rodriguez Bravo, Andrew Altieri, Julie L. Meyer

## Abstract

Global climate change impacts ocean communities through rising surface temperatures, ocean acidification, and deoxygenation. While the response of the coral holobiont to the first two effects has been relatively well studied, little is known about the response of the coral microbiome to deoxygenation. In this study, we investigated the response of the microbiome to hypoxia in two coral species that differ in their relative tolerance to hypoxia. We conducted *in situ* oxygen manipulations on a coral reef in Bahía Almirante, Panama, which has previously experienced episodes of low dissolved oxygen concentrations. Naïve coral colonies (previously unexposed to hypoxia) of massive starlet coral (*Siderastrea siderea*) and Lamarck’s sheet coral (*Agaricia lamarcki*) were transplanted to a reef and either enclosed in chambers that created hypoxic conditions or left at ambient oxygen levels. We collected samples of surface mucus and tissue after 48 hours of exposure and characterized the microbiome by sequencing 16S rRNA genes. We found that the microbiomes of the two coral species were distinct from one another and remained so after exhibiting similar shifts in microbiome composition in response to hypoxia. There was an increase in both abundance and number of taxa of anaerobic microbes after exposure to hypoxia. Some of these taxa may play beneficial roles in the coral holobiont by detoxifying the surrounding environment during hypoxic stress. This work describes the first characterization of the coral microbiome under hypoxia and is an initial step toward identifying potential beneficial bacteria for corals facing this environmental stressor.

**Importance:** Marine hypoxia is a threat for corals but has remained understudied in tropical regions where coral reefs are abundant. Deoxygenation on coral reefs will worsen with ongoing climate change, acidification, and eutrophication. We do not yet understand the response of the coral microbiome to hypoxia, and whether this reaction may have a beneficial or harmful role in the coral holobiont. To understand how the coral microbial community structure responds during hypoxic stress, we experimentally lowered the oxygen levels around corals in the field to observe changes in the composition of the coral microbiome. We documented the increase of anaerobic and pathogenic bacteria in the microbiomes of the massive starlet coral (*Siderastrea siderea*) and Lamarck’s sheet coral (*Agaricia lamarcki*) in 48 hours. This work provides fundamental knowledge of the microbial response in the coral holobiont during hypoxia and may provide insight to holobiont function during stress.

## INTRODUCTION

Marine deoxygenation is a devastating and global threat to oceanic and coastal ecosystems, with ecological, evolutionary, and social repercussions comparable to other major anthropogenic threats including warming and ocean acidification (1, 2). While previous work has established hypoxia as a widespread threat to temperate marine ecosystems (2–4), it has only recently garnered attention in tropical marine systems as a cause of mass mortality that reduces biodiversity and productivity (5). Many marine species globally are already in decline due to oxygen levels at or below critical oxygen thresholds (6), and decreased oxygen availability will likely be responsible for large shifts in ecosystem structures (7). Localized coastal hypoxia in tropical and subtropical waters has recently become a substantial threat to corals (8). Prolonged exposure to hypoxia can have adverse effects on coral health and resiliency including bleaching, disease, and mortality (5, 9–11).

Though prolonged exposure to hypoxia will ultimately lead to death, corals and other reef-associated organisms may have an innate tolerance to periodic deoxygenation (5, 6, 12–15). Corals undergo natural diel shifts in oxygen concentrations on their surface microenvironment (16–18). When sunlight is available in the photic zone during the day, *Symbiodiniaceae* oxygen production saturates the coral surface (16, 17). At night, coral holobiont respiration uses the free oxygen, creating a hypoxic microenvironment on the coral surface until sunlight triggers photosynthesis (16, 17). These diel changes in oxygen concentration can occur in the matter of minutes (18), yet the coral remains mostly undisturbed.

Corals may also experience some hypoxia tolerance during the periodic macroscale oxygen depletion that can occur naturally on reefs. These shifts in dissolved oxygen concentrations occur because of unusual weather patterns (19–21), reef geomorphology (19, 22–24), isolation of reefs during diel tidal cycles (22, 25), coral spawn slicks (20, 26), or other elements that reduce water column mixing and exchange with the open ocean (27). However, these natural occurrences of deoxygenation are exacerbated by eutrophication and climate change, intensifying the overall severity and duration of hypoxic events globally (1, 3, 8, 28, 29). With over 13% of the world’s coral reefs at an elevated risk for deoxygenation (5), understanding the response of and implementing mitigation strategies to coral reefs is critical.

The coral microbiome is a source of resilience for environmental stressors including warming (30, 31), and may play a similarly important role for hypoxia. Members of the microbiome fill a variety of functional roles within the coral host (9, 32–35), including nutrient cycling within the holobiont (33–35), nitrogen fixation (33, 34, 36), and pathogen resistance (33–35, 37). If there is flexibility of microbial species in response to dynamic oxygen conditions, this could contribute to the observed ability of coral hosts to withstand exposure to hypoxic conditions. Here, we experimentally induced hypoxic conditions with an *in situ* reef experiment to test how the microbiomes of the hypoxia-resistant massive starlet coral (*Siderastrea siderea*) (38) and the hypoxia-sensitive whitestar sheet coral (*Agaricia lamarcki*) (5, 38) responded to hypoxia.

## MATERIALS & METHODS

### Site description

Bahiá Almirante in Bocas del Toro, Panama is a large, semi-enclosed tropical embayment of 450 km^2^ (5) and is home to many shallow water (<25m) coral reefs (39, 40). This basin on the Caribbean coast shares many features with temperate estuaries that experience bouts of hypoxia, including reduced exchange with the open ocean, seasonal cycles of low wind energy and high temperatures, and a watershed delivering excess nutrients from agricultural run-off and untreated sewage (39, 41). Because of these conditions, Bahiá Almirante has experienced patches of hypoxic stress, with documented occurrences in 2010 and 2017 that caused extensive coral bleaching and necrosis in other marine invertebrates (5, 38). Due to these periodic hypoxic events, Bahiá Almirante and its coral reefs are ideal study sites for documenting coral health and resilience when exposed to low oxygen conditions. We chose massive starlet coral (*Siderastrea siderea*) and whitestar sheet coral (*Agaricia lamarcki*) as our study species because they are two of the predominant coral species in the region and exhibited strikingly different responses to prior hypoxia events, with *S. siderea* persisting at hypoxic sites (38), and *A. lamarcki* suffering near total mortality (5, 38).

### *In situ* oxygen manipulation

To test the response of coral microbiomes to hypoxic stress, we conducted a field experiment in which we manipulated oxygen with benthic incubation chambers. The experiment was conducted at Punta Caracol, in the vicinity of areas with documented mortality associated with hypoxia (Figure 1) (38, 42). Seven 60 x 60 cm plots were established and a miniDOT dissolved oxygen logger (Precision Measurement Engineering, Vista, CA) in each plot recorded oxygen concentration and temperature at 10-minute intervals. Four randomly selected plots were assigned to the hypoxia treatment, and the remaining three served as control plots (Figure 1). To create low oxygen conditions, 4-sided benthic incubation chambers were made of greenhouse-grade plastic. The chambers were open at the bottom, with 15-cm flanges that were tucked into the sediment to better isolate the water within. A submersible aquarium pump was placed in each chamber to homogenize the water column within.

**Figure 1:**
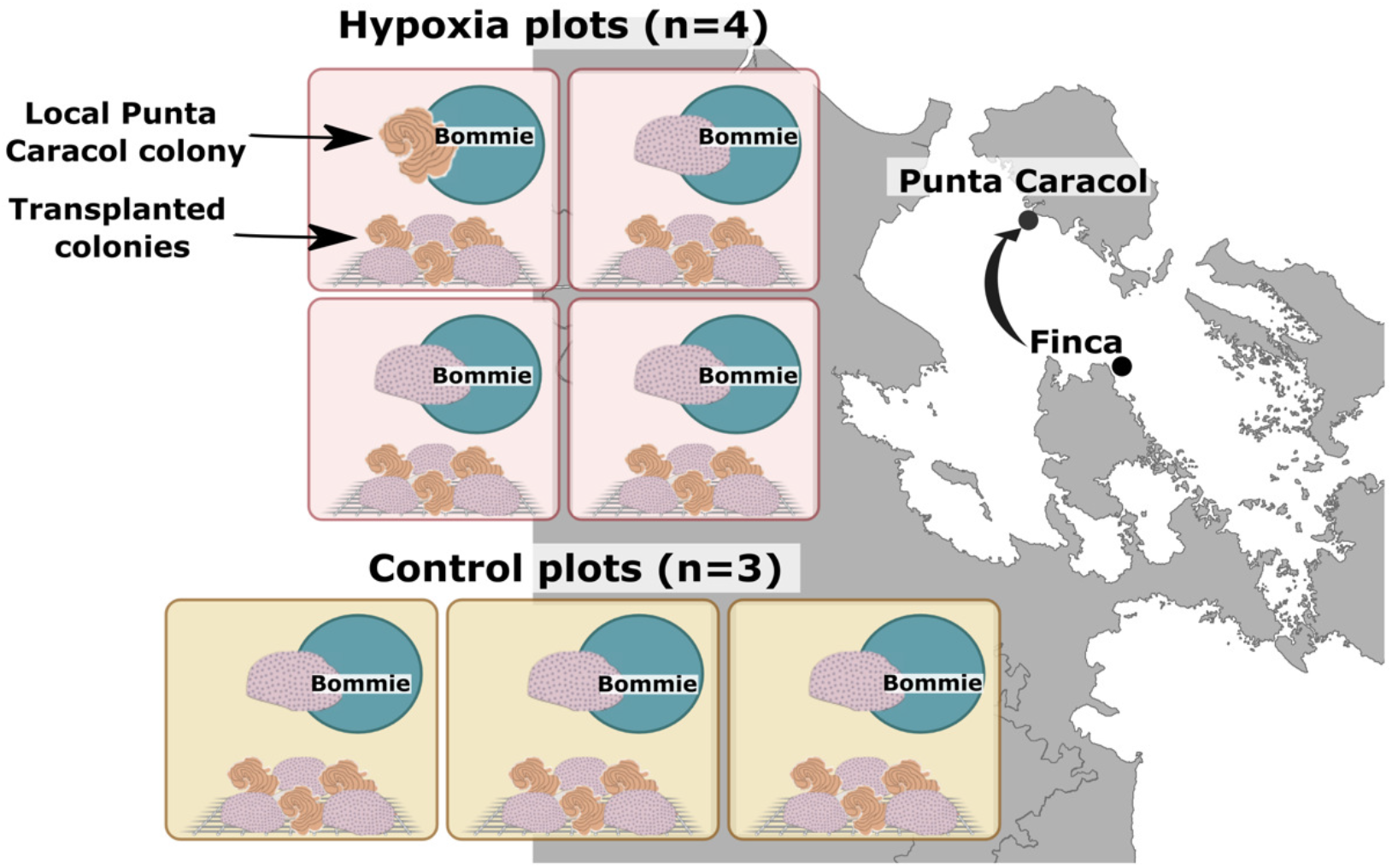
Map of experimental sites in Bahía Almirante, Bocas del Toro, Panama. Corals from Finca were transplanted to Punta Caracol for oxygen manipulation experiments (control plots, hypoxic plots). Each of the seven plots contained a mixed species bommie with a local Punta Caracol colony attached. Three transplanted *S. siderea* colonies and *A. lamarcki* were also placed in each plot by fastening the colonies to a mesh rack.

Colonies of *A. lamarcki* and *S. siderea* (7 - 12 cm diameter) were collected at the Finca site from a depth of 5-10 m for transplantation to the experimental plots. Colonies were collected at least 2m apart and likely represented independent genotypes. Coral colonies were transported in aerated seawater to Punta Caracol where they were randomly assigned to experimental plots. Each plot contained a local Punta Caracol bommie with a representative reef community that contained a mix of corals, sponges, and other benthic organisms that included either a *S. siderea* or *A. lamarcki* colony. We transplanted three *S. siderea* and three *A. lamarcki* colonies to each plot by fastening the colonies to a mesh rack next to the bommie (Figure 1). The experimental oxygen manipulation was conducted for 48 hours, at which time the coral surface microbiome was sampled.

### Coral microbiome sampling

In addition to coral colonies in the experimental plots, three colonies of *S. siderea* were sampled from Tierra Oscura where hypoxia has been previously documented and three colonies each of *A. lamarcki* and *S. siderea* were sampled from Finca where hypoxia has not been documented (Figure S1) (38, 42, 43). Coral mucus/tissue samples were collected by agitation and suction of the coral surface with individual sterile needleless syringes. Syringes were transported in a cooler with ice to the lab, and mucus was allowed to settle in the syringes before expelling into a 2-ml cryovial with RNALater (Ambion, Austin, TX). Preserved samples were frozen until further processing at the University of Florida.

### V4 amplicon library preparation

Extraction of genomic DNA was performed with a DNeasy Powersoil kit (Qiagen, Germantown, MD) according to manufacturer instructions. The V4 region of the 16S rRNA gene was amplified in triplicate for each sample using the 515F (44) and 806RB (45) Earth Microbiome primers and thermocycler protocol (46) in 25-μl reactions containing Phusion High-fidelity Master Mix (New England Biolabs, Ipswich, MA), 0.25μM of each primer, 3% dimethyl sulfoxide (as recommended by the manufacturer of the polymerase), and 2 μl of DNA template. Triplicate reactions were consolidated and cleaned with a MinElute PCR purification kit (Qiagen) and quantified with a DS-11 FX+ spectrophotometer (DeNovix, Wilmington, DE). One DNA extraction kit blank without the addition of any starting coral biomass was produced alongside regular DNA extractions, and then amplified and sequenced using a unique barcode. One final pool containing 240 ng of each amplicon library was submitted to the University of Florida Interdisciplinary Center for Biotechnology Research (RRID:SCR_019152) for sequencing on an Illumina MiSeq with the 2×150bp v.2 cycle format.

### Analysis of V4 amplicon libraries

Quality filtering, error estimation, merging of reads, dereplication, removal of chimeras, and selection of amplicon sequence variants (ASVs) were performed with DADA2 v. 1.18.0 (47) in RStudio v. 1.1.456 with R v. 4.0.4. Taxonomy was assigned to ASVs using the SILVA small subunit rRNA database v. 132 (48). The ASV and taxonomy tables were imported into phyloseq v. 1.34.0 (49) for analysis and visualization of microbial community structure. ASVs assigned as chloroplast, mitochondria, or eukaryote were removed from further analysis. Variation in community composition was determined using the Aitchison distance of centered log-ratio transformed, zero-replaced read counts using CoDaSeq v. 0.99.6 (50) and visualized with principal component analysis (PCA). Principal component analysis of the Aitchison distance was performed with the package prcomp in R and plotted with ggplot2 v. 3.3.3 (51). Permutational Multivariate Analysis of Variance (PERMANOVA) with vegan v. 2.5-7 (52) was used to test for differences in community structure by treatment and coral species. For clarity, the nine coral microbiome samples collected at Tierra Oscura and Finca that were not part of the experimental plots were only included in the Analysis of Compositions of Microbiomes (ANCOM) figures, as they did not provide sufficient statistical power for additional analyses. ANCOM (53) was used to identify microbial families that were differentially abundant across treatments, using an ANOVA significance level of 0.05 and removing families with zero counts in 90% or more of samples. Only families detected in at least 70% of samples were reported. Finally, indicspecies v. 1.7.9 (54) was used to identify differentially abundant ASVs amongst treatment types. The complete set of R scripts and metadata are available at github.com/meyermicrobiolab/Panama_Hypoxia.

## RESULTS

### Experimental deoxygenation

Dissolved oxygen (DO) concentrations (mg/L) in the control plots ranged from 4.29 mg/L - 6 mg/L throughout the experimental period, while DO concentrations in hypoxia chambers steadily decreased (Figure 2A). In the chamber associated with MiniDOT logger 3, DO concentrations decreased drastically starting at hour 8, and reached levels <0.1 mg/L around hour 15 of the experiment (Figure 2A). At hour 15, the other hypoxia chambers were at 2-3 mg/L, while our open chambers were at 5.5-6 mg/L, and the oxygen concentrations in the chambers continued to decline thereafter. Because of this extreme decrease, we observed *in situ* that corals within chamber 3 experienced severe bleaching. Over the course of 48 hours, water temperature ranged from 29.42°C − 30.08°C in the Punta Caracol experimental plots (Figure S2).

**Figure 2:**
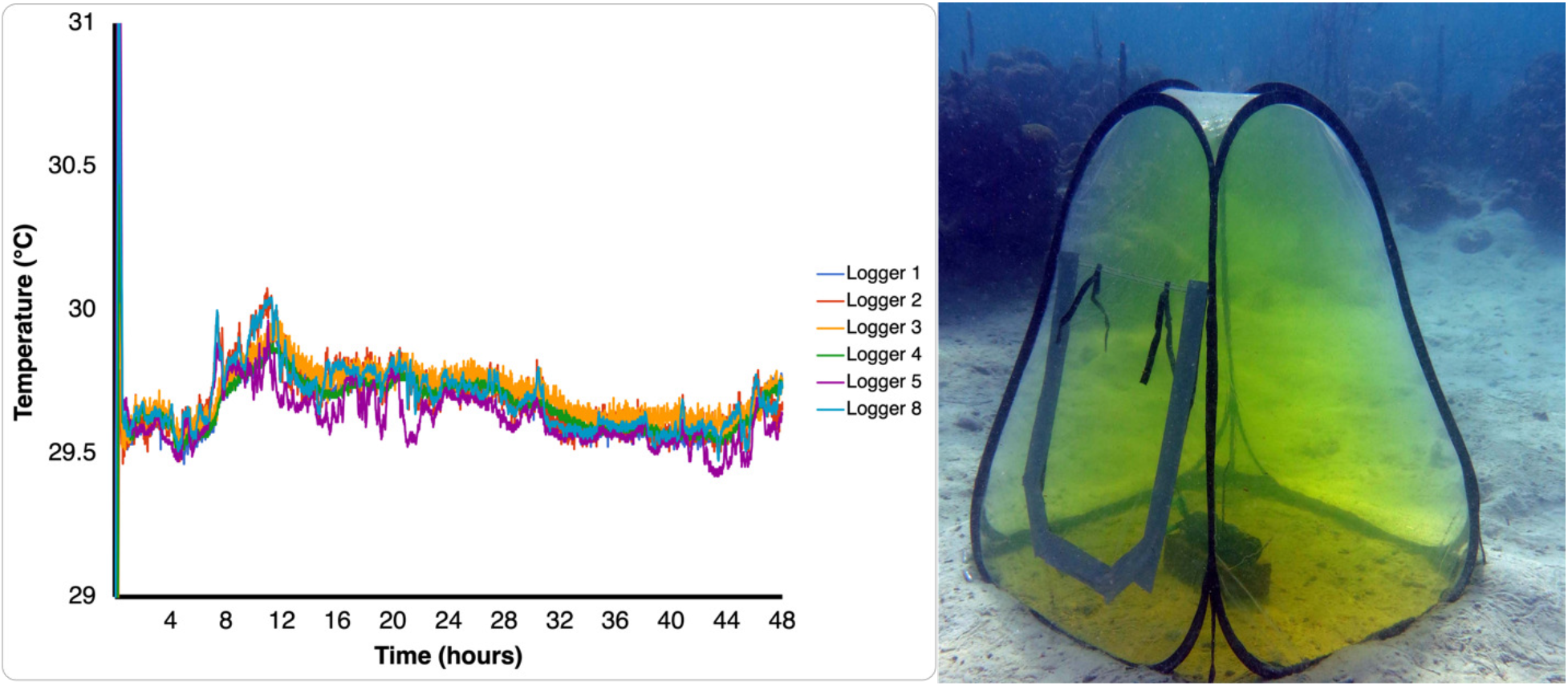
(A) Dissolved oxygen concentrations (mg/L) in the hypoxic and control plots over 48 hours. Tent 3 became hypoxic rapidly and stayed hypoxic for the duration of the experiment. (B) An example of the greenhouse chamber used to simulate natural hypoxia in the marine environment. Fluorescein dye was used in trials to ensure the chambers could be secured with minimal flow-through and leaks.

### Microbial community characterization

Microbial communities were characterized for a total of 56 coral mucus samples from *Agaricia lamarcki* and *Siderastrea siderea* collected from three different sites in May 2019 (Figure S1, Table S1). After quality-filtering and joining, an average of 56,660 sequencing reads (11,273–116,996) per coral sample were used in the analysis (Table S1). A total of 19 archaeal ASVs and 860 bacterial ASVs were detected. One control sample from the extraction kit was also sequenced, and after quality-filtering and joining it had 22,860 reads, which were classified as 78 bacterial ASVs (Table S2). Sequencing reads with primers and adapters removed are available at NCBI’s Sequence Read Archive under BioProject PRJNA641080.

Overall, microbial community structure in the experimental plots differed by coral species, although the effect size was small (PERMANOVA, P=0.001, R^2^=0.08, Figure 3).

**Figure 3:**
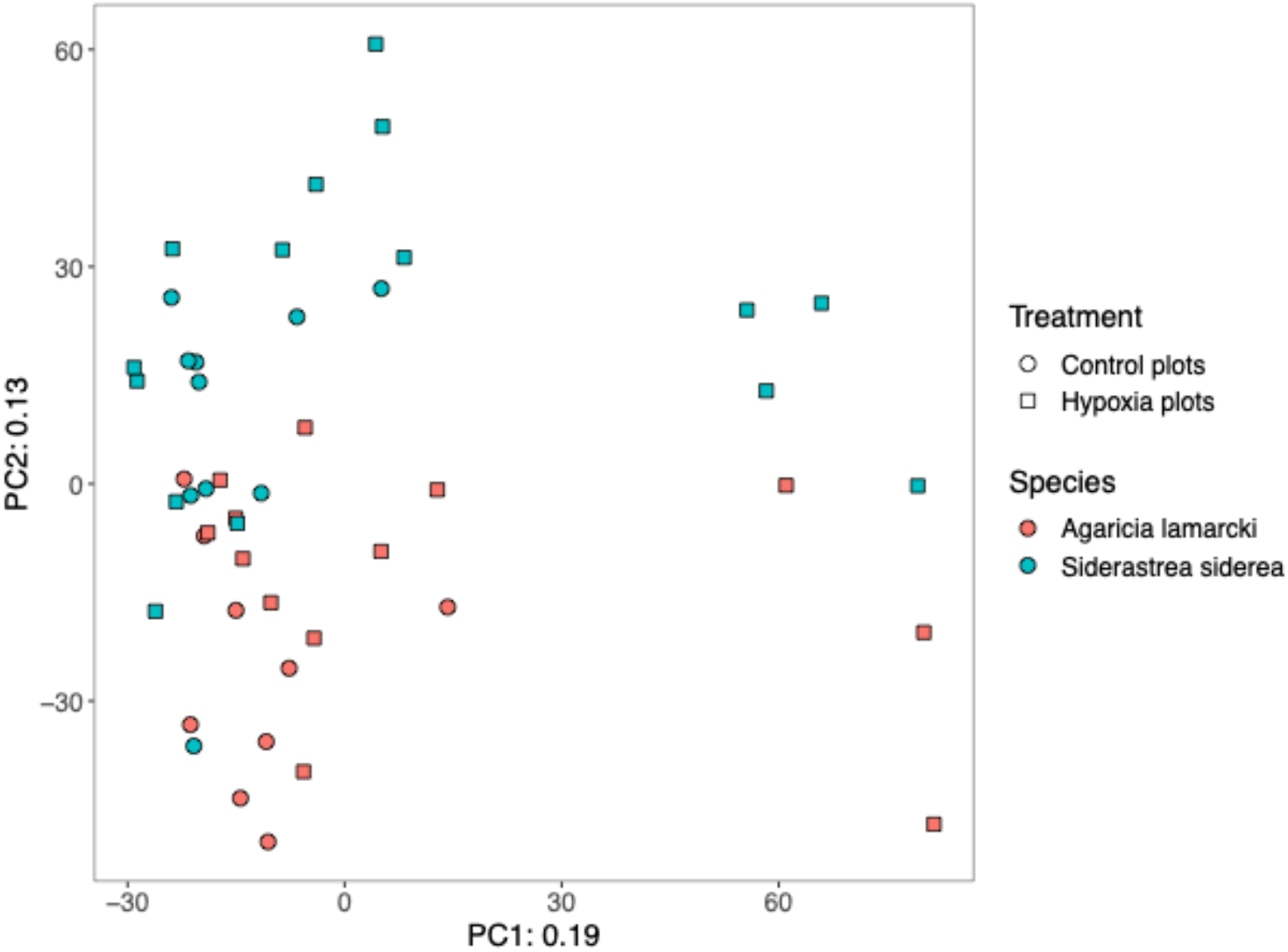
Principal component analysis of microbial community structure in corals in the control plots and corals in the hypoxia plots.

Additionally, microbial community structure differed among corals in the control plots and the hypoxia plots, although the effect size was small (PERMANOVA, P=0.001, R^2^=0.06, Figure 3). The interaction between coral species and treatment was not significant (PERMANOVA, P>0.05, R^2^=0.02). Additionally, there was no significant difference in coral microbial community structure among the six unmanipulated *S. siderea* sampled in Tierra Oscura and Finca (ANOSIM, P>0.05, R^2^=0.63).

*Alphaproteobacteria*, *Gammaproteobacteria*, and *Bacteroidia* were commonly detected in all samples, regardless of treatment and species (Figure 4), consistent with previous studies of coral microbiomes (55). All ASVs classified only as Bacteria (n=22) were searched with BLASTn and sequences labeled as mitochondria by NCBI were removed from the data set. The most abundant ASV classified only as Bacteria in both species (Figure 4) was 87% similar to an uncultivated bacterial sequence associated with the cold-water coral *Lophelia pertusa* sampled in Norway (GenBank Accession AM911366) (56) based on BLASTn searches. Additionally, the most abundant ASV classified only to class *Gammaproteobacteria* was 98% similar to an uncultivated Caribbean coral-associated bacterium (GenBank Accession KU243233) (57). The most abundant ASV classified only to phyla *Proteobacteria* in *S. siderea* (Figure 4B) was 92% similar to an uncultivated *Deltaproteobacteria* associated with the coral *Pavona cactus* originating from the Red Sea (GenBank Accession EU847601) (58).

**Figure 4:**
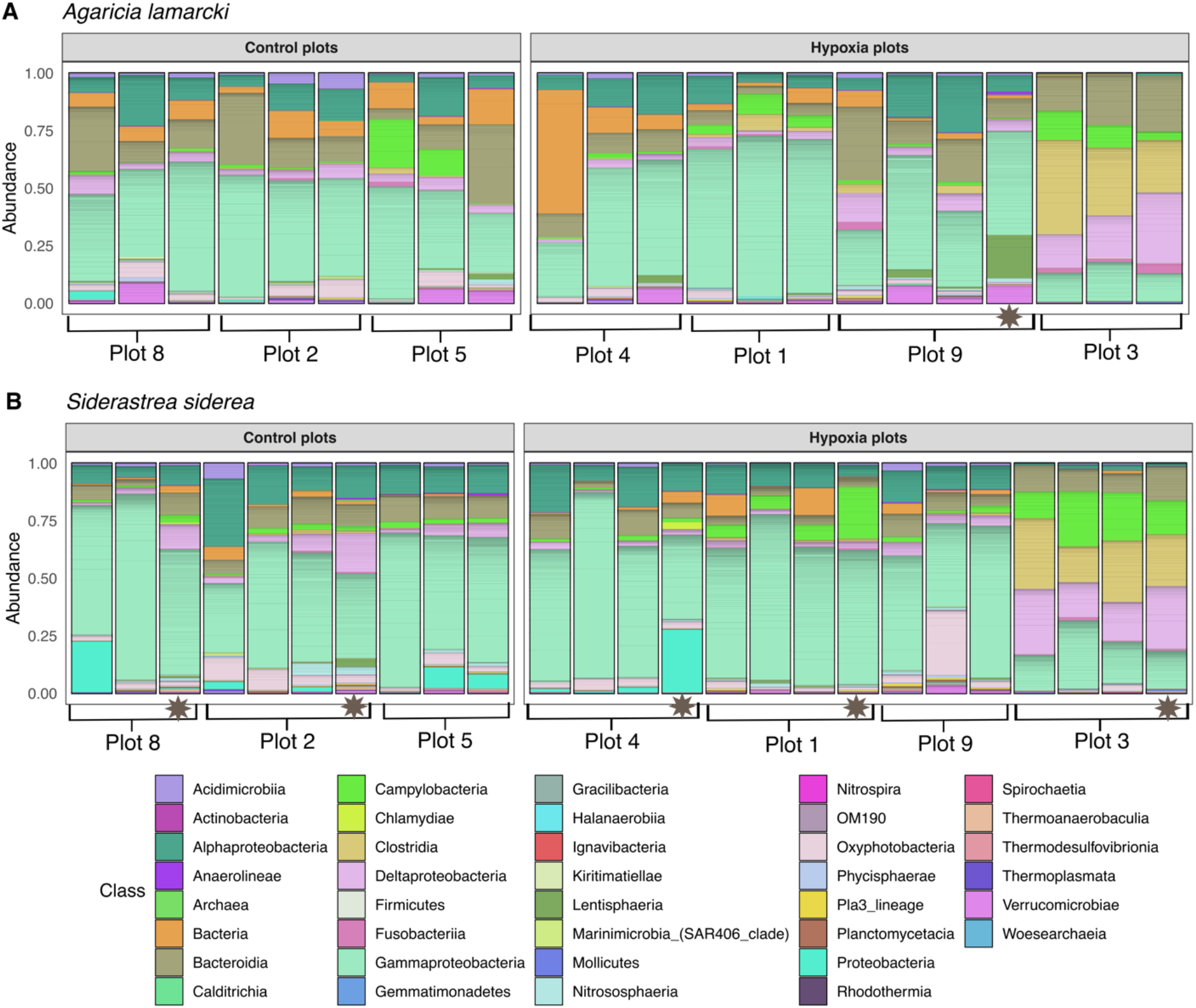
Relative abundance of amplicon sequence variants, colored by class, in corals in the control plots, and corals in the hypoxia plots for *Agaricia lamarcki* (A) and *Siderastrea siderea* (B). Gray stars indicate local Punta Caracol coral colonies in the plots.

Differences among treatments in the microbial community structure were primarily driven by 14 differentially abundant families (Figure 5). These families were detected in at least 70% of the samples and were significantly different (ANOVA, p=0.05) among unmanipulated corals from Tierra Oscura and Finca, control plots, and hypoxic plots (Figure 5). The largest differences among treatment types were observed in families *Desulfovibrionaceae*, *Nitrincolaceae*, *Clostridiales Family XII*, and *Midichloriaceae*. The relative abundances of *Desulfovibrionaceae*, *Nitrincolaceae*, *Clostridiales Family XII* were higher in the hypoxia treatment, whereas family *Midichloriaceae* was highest in the unmanipulated corals (Figure 5). *Clostridiales Family XII* was more abundant in corals exposed to hypoxia and less abundant in unmanipulated and control plot corals.

**Figure 5:**
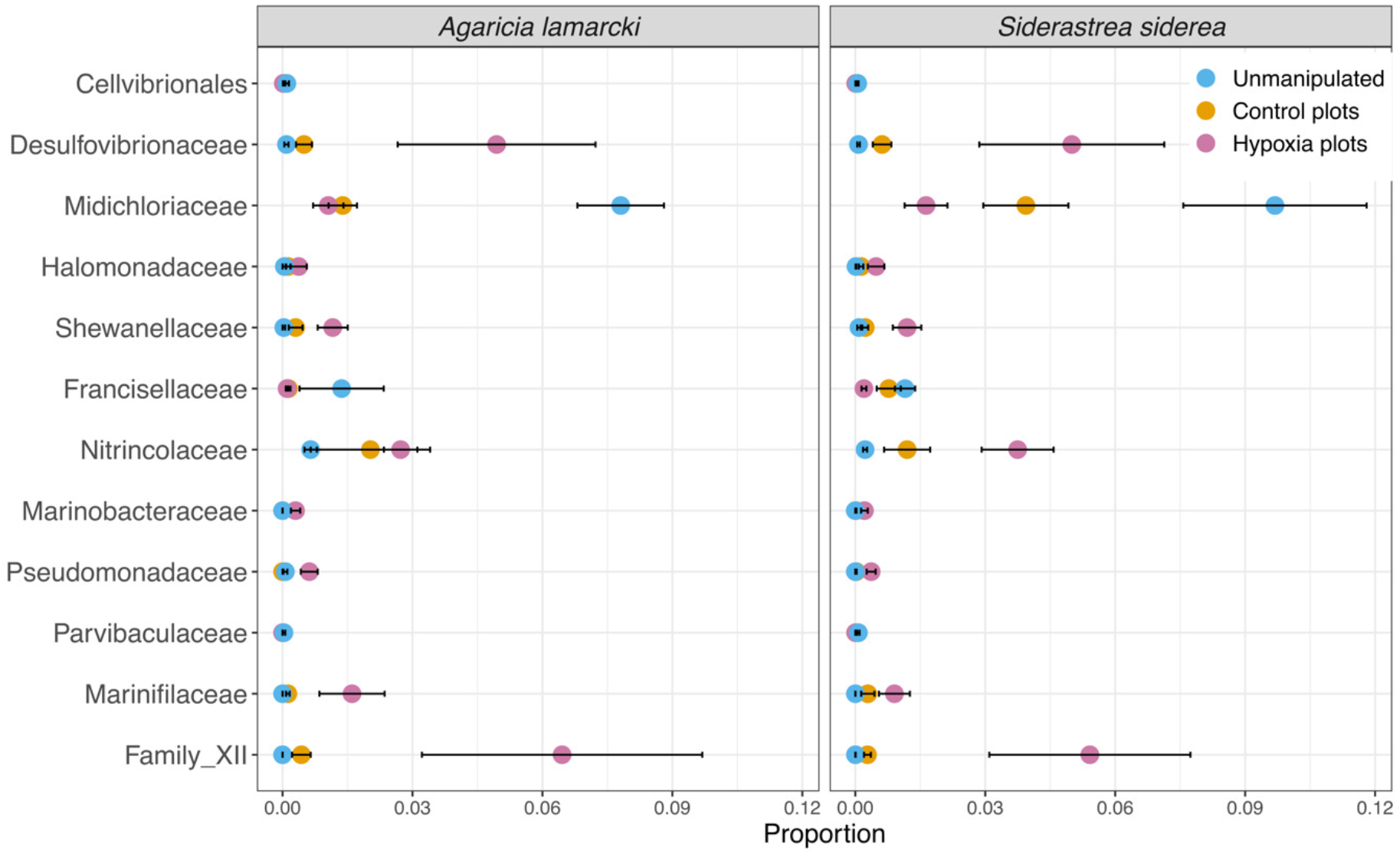
Relative abundance of 14 microbial families that were differentially abundant across treatment types: unmanipulated corals from Finca and Tierra Oscura, corals in the control plots, and corals in the hypoxic plots. Points represent the average relative abundance and error bars depict the standard error from analysis of all 56 coral samples.

The hypoxic chamber 3 experienced a sudden and dramatic drop in dissolved oxygen concentrations and was completely hypoxic for 36 hours. This was associated with the largest magnitude response of the microbiome relative to the other plots. The seven microbial communities grouped on the right side of the PCA (Figure 3) were from corals exposed to extremely low dissolved oxygen concentrations in chamber 3 (Figure 2A) that ultimately bleached. The corals in this chamber included three colonies of *A. lamarcki* and four colonies of *S. siderea*, one of which was a local Punta Caracol *S. siderea* colony. Microbial community structure varied more by chamber (PERMANOVA, P=0.001, R^2^=0.37) than by either species or treatment. Microbial community shifts in chamber 3 with 36 hours of hypoxia exposure are clearly seen in the relative abundances of microbial classes in both coral species (Figure 4). Microbial communities in this plot experienced a dramatic increase in *Clostridia*, *Deltaproteobacteria*, and *Campylobacteria*, groups that are composed of numerous anaerobes (Figure 4).

Differences among the plots were primarily driven by 41 differentially abundant bacterial families. Those that were detected in higher abundances in both coral species from chamber 3, which was the most hypoxic plot, include *Arcobacteraceae*, *Prolixibacteraceae*, *Marinilabiliaceae*, *Desulfobacteraceae*, *Bacteroidales*, *Peptostreptococcaceae*, *Desulfovibrionaceae*, *Marinifilaceae*, and *Clostridiales Family XII* (Figure S3). The relative abundances of *Midichloriaceae* were lowest in chamber 3, as were unclassified *Gammaproteobacteria* and *Proteobacteria* families (Figure S3). Families *Colwelliaceae* and *Vibrionaceae* were detected in higher abundances in hypoxic chamber 1. Both coral species had several families in common that similarly showed responses of large magnitude to hypoxia, including *Arcobacteraceae*, *Desulfovibrionaceae*, and *Clostridiales Family XII* (Figure 6).

**Figure 6:**
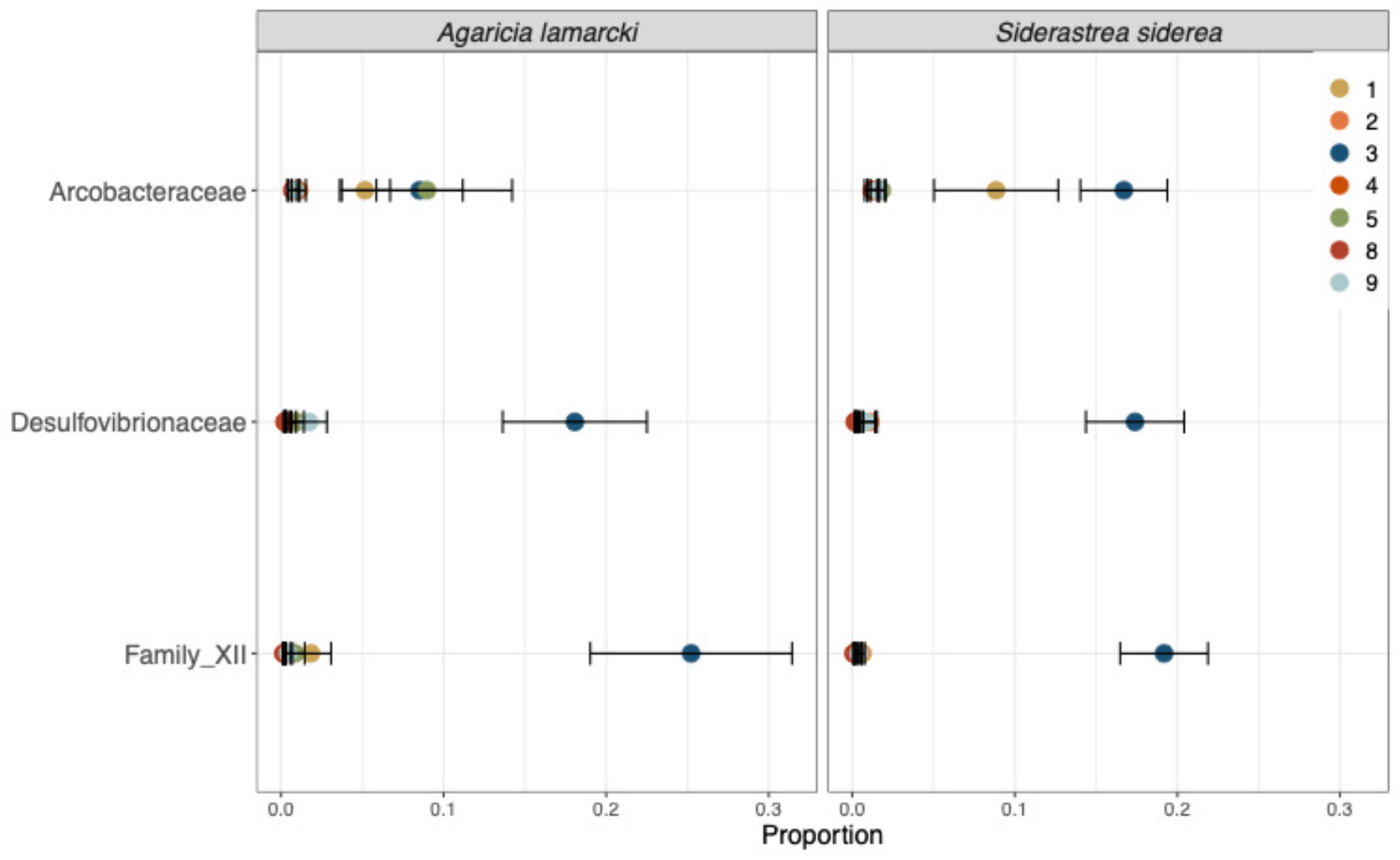
Relative abundance of 3 families that were differentially abundant across chambers. Colored points represent the average relative abundance of the families in each plot and error bars depict the standard error from analysis of 47 coral samples. Families *Arcobacteraceae*, *Clostridiales Family XII*, and *Desulfovibrionaceae* increased significantly in corals that experienced hypoxia for the longest (36 hours).

To see if differentially abundant families were driven by particular ASVs, an indicator species analysis was performed on all samples. Of the 878 ASVs tested, 144 ASVs were considered indicator species for hypoxia, but only four ASVs had a correlation statistic ≥ 0.50 (Figure S4). These include an *Alteromonas* ASV, a *Neptuniibacter* ASV, an *Aestuariicella* ASV, and a *Marinobacter* ASV (Table S3). We performed another indicator species analysis for each coral species individually to determine if any ASVs shifted in both coral species when exposed to hypoxic stress, and we detected no indicator ASVs common to both species. Therefore, although the two coral species exhibited similarity in the identity of microbial families that were differentially abundant and in the directionality of their shifts in abundance, these patterns were not driven by individual taxa common to both *A. lamarcki* or *S. siderea*.

## DISCUSSION

We observed a shift in the microbial communities of coral *A. lamarcki* and *S. siderea* under experimental deoxygenation in less than 48 hours. This shift was deterministic rather than stochastic as the microbial communities in both coral species responded in similar ways.

Hypoxic conditions resulted in an increase of anaerobic and potentially pathogenic bacteria in the classes *Deltaproteobacteria*, *Campylobacteria*, and *Clostridia* in the microbiome of both *A. lamarcki* and *S. siderea*. This is most apparent in corals that experienced the most severe hypoxia associated with plot 3. Moreover, both coral species exhibited changes of similar magnitude in the relative abundances of many families, most notably *Arcobacteraceae*, *Desulfovibrionaceae*, *Clostridiales Family XII*, *Nitrincolaceae*, and *Midichloriaceae*. Though a difference in microbial communities between oxygen treatments for both species was statistically different, the effect size of that difference was relatively small. In contrast, previous work has shown stochastic shifts in the microbiome in response to other environmental pressures. For example, stressors including nutrient pollution, overfishing, and thermal stress on reefs were correlated with an increase in the dispersion of beta diversity in the coral microbiome (60). Our results suggest that there may be some host regulation of the microbiome in response to hypoxic stress, as the microbiome changed in a similar fashion between the two species. Instead of having stochastic responses in the microbiome, these corals may curate the members of their microbial community to better deal with the stress of deoxygenation (61). Examining the functional role of these members may explain the uniformity of the microbiome across both coral species in response to hypoxia.

### Functional significance of microbiome shifts

Under experimentally induced hypoxia, we documented an increase in *Deltaproteobacteria*, specifically the family *Desulfovibrionaceae*. *Deltaproteobacteria* are known for their role as sulfate-reducing microorganisms (SRM) (62, 63). In marine ecosystems, *Deltaproteobacteria* are mainly found in sediment, where they are the predominant SRMs in terms of abundance and activity (64). *Desulfovibrionaceae*, a well-known family within *Deltaproteobacteria*, includes numerous sulfate-reducing species which produce hydrogen sulfide that can degrade coral health. Members of this family have been implicated in Black Band Disease as a producer of sulfide (65, 66). Further, *Desulfovibrionaceae* were detected in corals infected with stony coral tissue loss disease (SCTLD), and the genera *Desulfovibrio* and *Halodesulfovibrio* have been recently described as bioindicators of the disease (67, 68). *Deltaproteobacteria* in the coral microbiome are likely producing sulfide and playing an antagonistic role and may contribute to increased coral disease prevalence associated with reef hypoxia, but the definitive role of this class in the coral microbiome remains to be confirmed, particularly under environmental stressors like hypoxia.

We also documented an increase in the class *Campylobacteria* during experimental deoxygenation in the coral microbiome. Microbes within this taxonomic group, and many species of *Epsilonbacterota* in particular, play important roles in carbon, nitrogen, and sulfur cycling, especially in symbiosis with their host (69, 70). *Epsilonbacterota* thrive in anaerobic or microaerobic environments rich with sulfur (69), including hydrothermal vents (69) and sediments associated with seagrass roots (71). On corals experiencing hypoxia, members of *Campylobacteria* may alleviate stress by oxidizing some of the toxic sulfides produced by microbial respiration including *Deltaproteobacteria* in the holobiont. The increase in sulfur-oxidizing *Campylobacteria* during hypoxia may therefore be a form of rapid adaptation to this stressor, conferring resilience to deoxygenation stress for corals. For instance, family *Arcobacteraceae*, which were enriched under the most extreme low oxygen conditions here, are known for the sulfide-oxidizing capabilities (72, 73), producing both sulfate and filamentous sulfur (73). *Arcobacteraceae* are associated with changes in the coral holobiont under stress conditions, growing rapidly in the microbiome in thermally stressed corals (74) and corals living in polluted waters (75). While members of this group have also been associated with coral diseases, such as white syndrome (76), brown band disease (76), white plague disease (77), and stony coral tissue loss disease (68), an increase in *Arcobacteraceae* during hypoxic stress may play a beneficial role in the coral microbiome.

*Clostridia*, including *Clostridiales Family XII*, also increased in abundance on both species of coral host in response to deoxygenation. This change is especially prominent in chamber 3, where hypoxia was most severe and sustained. *Clostridia* is a large polyphyletic class of obligate and facultative anaerobes known for producing the highest number of toxins of any bacterial group and causing severe disease in humans and animals (78). However, the role of *Clostridia* in coral remains ambiguous. Most Gram-positive sulfate-reducing bacteria belong to the class *Clostridia*, so these taxa may play a similar role to the *Deltaproteobacteria* in the coral holobiont (79). Further, corals that harbor higher abundances of *Clostridia* ASVs are more often associated with disease (80). For example, *Clostridiales* ASVs are enriched in the surface mucus layer and tissue near stony coral tissue loss disease (SCTLD) lesions (59, 68, 81) and black band disease mats (82, 83). An increase of *Clostridia* has also been documented in the microbiome when corals are exposed to thermal stress (84). Generally, higher abundances of *Clostridia* in the coral microbiome are often associated with host stress. In our study, members of *Clostridia* are likely playing an antagonistic role in the coral holobiont as sulfide producers (79) or as opportunistic pathogens as oxygen levels decline (80). However, *Clostridia* remains unsubstantiated as the causative agent of any coral disease, and it may simply respond opportunistically to stress-associated changes in the holobiont.

Family *Nitrincolaceae*, belonging to class *Gammaproteobacteria*, was more abundant in corals exposed to hypoxia. This increase in *Nitrincolaceae* is consistent with observations in the microbial community in the water column above a reef during the 2017 hypoxic event in Bahía Almirante when *Nitrincolaceae* was found only in hypoxic water samples from that event, and not in oxygenated water samples at that site following the event or at a reference site (38). The genus *Amphritea* within the family *Nitrincolaceae* is considered a bioindicator for stony coral tissue loss disease in *S. siderea* (67). Species within this family have genes for nitrite reductase, nitric oxide reductase, and nitrous oxide reductase (85, 86). As such, members of *Nitrincolaceae* have the potential to produce nitrate (NO_3_), nitrous oxide (N_2_O), and dinitrogen (N_2_). The denitrification of bioavailable nitrogen to nitrogen gas in this system may aid in mitigating the eutrophication that usually precedes hypoxia (29). Taxa within this family have also been described as following short term “feast and famine” dynamics of nutrient uptake and are aggressive heterotrophs (86). During seasonal transitions in the Southern Ocean, *Nitrincolaceae* rapidly take up nutrients from phytoplankton-derived organic matter and iron (86). In hypoxic conditions on coral reefs, it is possible that our observed increase in *Nitrincolaceae* signified their role as opportunistic heterotrophs. Their increase in the holobiont may be due to coral tissue decay, as death of both coral and associated *Symbiodiniaceae* may supply the bacteria with the organic matter and iron they need to thrive in this environment.

Family *Midichloriaceae* (order *Rickettsiales*) decreased in all corals associated with hypoxic conditions, including those in chamber 3. *Rickettsiales* are obligate intracellular bacteria of eukaryotes and include well known zoonotic pathogens (87). Though previously implicated in white band disease (88, 89), many recent studies have detected the *Rickettsiales* genus MD3-55 (*Candidatus* Aquarickettsia rowherii) as an abundant member of the apparently healthy *Acropora cervicornis* microbiome in the Cayman Islands (90), the Florida Keys (91–93), and Panama (94, 95). *Rickettsiales* have also been found in low abundances on six healthy coral species sampled in the Bocas del Toro region of Panama (95). In our study, family *Midichloriaceae* were detected at lower relative abundances under hypoxic conditions. This may be due to some tissue loss in corals that experienced severe hypoxia in chamber 3 and indicate that *Rickettsiales* has a dependence or preference for healthy corals. Though their role in the coral microbiome remains unclear, our study provides further evidence that *Rickettsiales* is a constituent of healthy holobiont that declines in abundance with stress.

### Holobiont response to hypoxic stress

Differences in hypoxia tolerance thresholds among coral species may be due to regime of hypoxia exposure, host stress responses, or microbial function. Environmental history can also affect the survival of coral during subsequent exposures to low oxygen (100). Previous work has demonstrated that coral species vary in their susceptibility to hypoxia (5, 96–99). For example, *Acropora cervicornis* suffered tissue loss and mortality within a day of exposure to hypoxia in lab experiments, whereas *Orbicella faveolata* was unaffected after 11 days of continuous hypoxia exposure (96). *Stephanocoenia intersepta* from Bahiá Almirante exhibited a threefold greater hypoxia tolerance than *A. lamarcki* in lab-based experiments (5). Further, following a deoxygenation event in Morrocoy National Park, Venezuela, *Acropora* and some *Montastrea* colonies exhibited bleaching, while *S. siderea*, *Porites astreoides,* and *P. porites* did not suffer any damage (97). These data follow a trend: plating and branching corals typically have a higher mortality rate than massive and encrusting corals under hypoxic conditions (21, 26, 97, 98, 100). These differences in hypoxia tolerance have been observed in prior studies done in Bahiá Almirante, which record *Agaricia* species as hypoxia sensitive (5, 38), and *S. siderea* as hypoxia resilient (38).

In addition to innate resilience that appears to vary with morphology, transcriptomic analysis has revealed that corals possess a complete and active hypoxia-inducible factor (HIF)-mediated hypoxia response system (HRS) that confers some hypoxia resilience (99). The effectiveness of this hypoxia response system can differ between coral species. For example, *Acropora tenuis* was more resistant to hypoxic stress when compared to *Acropora selago*. *A. tenuis* exhibited bleaching resistance and showed a strong inducibility of HIF genes in response to hypoxic stress. In contrast, *A. selago* exhibited a bleaching phenotypic response and was accompanied by lower gene expression of the hypoxia-inducible factor (HIF)-mediated hypoxia response system (99). Therefore, differences in coral response to hypoxia are in part due to the effectiveness of their HIF-HRSs.

Though historic exposure and the HIF-HRS each contribute to host survival, it is likely a synergistic effect between historic exposure, the HIF-HRS, and the coral microbiome that confer the most resilience to the holobiont during hypoxia. Past research has demonstrated that corals may shuffle members of their holobiont to bring about the selection of a more advantageous microbiome in response to environmental stressors (33, 101, 102). This microbial shuffling may act as a form of rapid adaptation to changing environmental conditions rather than mutation and natural selection (61). In our results, we observed a rapid shift in the community composition of the microbiome in response to hypoxia associated with the survival of corals through a period of intense deoxygenation stress, supporting the idea of a flexible microbiome conferring adaptation. We presume that some microbial taxa that increased in abundance with hypoxia may play a role in host resilience by eliminating toxic natural products around the microenvironment of the coral or by filling some metabolic needs during stress. This appears to be a common overall strategy across coral species that has developed in response to the selective pressure of hypoxia given that we observed it across two species that are distantly related taxonomically and are at opposite ends of the spectrum with regards to hypoxia tolerance. However, the exact ASV constituents that contributed to the shifts at the family level differed between the corals, suggesting different co-evolutionary pathways which may contribute to the difference in hypoxia tolerance of the coral hosts.

### Conclusions

Marine deoxygenation will worsen with continued climate change, and with its potential to degrade coral reefs it is essential to understand patterns of resilience revealed in the microbiome. Given the results of this study, we suspect that increased abundances in some microbial taxa with hypoxia may play a role in host resilience by detoxifying the microenvironment around the coral host, such as *Campylobacteria* (*Arcobacteraceae*). Other taxa, such as *Midichloriaceae* and *Clostridiales Family XII*, have more ambiguous roles in the coral microbiome, though their shifts in response to hypoxia warrant further investigation. We hypothesize that enhancement of these anaerobes, facultative anaerobes, or microaerophiles in the microbiome fill necessary and diverse metabolic functions in the holobiont while simultaneously indicating deoxygenation. Future studies that examine the functional roles of the coral microbiome through metagenomic or metatranscriptomic analyses can further advance our understanding by testing these hypotheses as to how the microbiome can mitigate the degradation of coral reefs under hypoxic conditions.

### Data availability

Sequences are available on the NCBI Sequence Read Archive (SRA) BioProject PRJNA641080 under the accession numbers: SAMN15298019-SAMN15298075.

## ACKNOWLEDGMENTS

We thank the team at the Smithsonian Tropical Research Institute in Bocas del Toro, Panama for their assistance in field monitoring and work. This research was supported by University of Florida start-up funds to AHA and JLM, and NSF grant OCE-2048914 to AHA and JLM.

